# Inhibition of *Candida albicans* virulence factor by cyclic dipeptides derived from *Aeromonas veronii* V03

**DOI:** 10.1101/2024.10.24.619162

**Authors:** Jinendiran Sekar, Gokul Parasuraman, Dhanasekaran Dharumadurai, B.S. Dileep Kumar, Sivakumar Natesan

## Abstract

*Candida albicans* is the most common human fungal pathogen with high mortality rates and limited antifungal treatments. Inhibition of *C. albicans* pathogenesis by targeting virulence factors provides a promising strategy for the development of novel antifungal drugs and overcoming drug resistance. In this study, four structurally different cyclic dipeptides (or diketopiperazine(DKP) were isolated and identified as cyclo(L-Pro-L-Leu), cyclo(L-Pro-L-Val), cyclo(D-Pro-L-Phe), and cyclo(L-Pro-D-Tyr) from *Aeromonas veronii* V03 and their antimicrobial potentials were evaluated. Results revealed that identified DKPs exhibited antibacterial activity against bacterial pathogens, such as *Staphylococcus aureus, Proteus mirabilis*, *Pseudomonas aeruginosa,* and *Aeromonas hydrophila*. Importantly, cyclo(D-Pro-L-Phe) lacking hydroxyl groups showed potent inhibitory effects against *C. albicans* and non-albicans species with low concentrations. Moreover, identified DKPs inhibited the virulence traits of *C. albicans,* including yeast-to-hyphae transition, secreted hydrolases (aspartic proteases and phospholipase) and biofilm formation in a dose-dependent manner. Collectively, our findings suggest that cyclic dipeptides from DKPs derived from *A. veronii* V03 could potentially be developed as antivirulence agents against *C. albicans* infection.

## 1. Introduction

According to World Health Organization reports, the prevalence of infectious diseases caused by both Gram-positive and Gram-negative bacterial and fungal pathogens is considered a major threat to human health[1, 2]. For instance, *Candida albicans*, an opportunistic fungus, and *Staphylococcus aureus, Escherichia coli*, *Proteus mirabilis*, and *Pseudomonas aeruginosa* are ubiquitous bacterial pathogens and are among the top causes of serious nosocomial infections and invasive disease[1–4]. While these strains can cause significant morbidity and mortality on their own, they are often co-isolated from various colonization and infection niches and are correlated with more severe disease states and higher mortality rates, even with current therapeutic interventions[1, 2, 5, 6]. Emerging multidrug resistance among bacterial and fungal pathogens has necessitated the development of alternative approaches to combat resistant-associated infection than currently used treatment regimens[2, 6, 7]. However, recent studies have shown that inhibition of virulence factors may restrict fungal growth and is less likely to result in the acquisition of drug resistance[8–13]. Understanding the environmental factors that influence the switch between commensal and pathogenic states is crucial for the development of antivirulence therapies[8, 11]. It is well described *C. albicans* can undergo a morphological transition from the yeast form to filamentous hyphae is an important virulence trait and is associated with secreted hydrolases and candidalysin, as well as cell adhesion that promote tissue invasion and pathogenicity[14–16]. Yeast to-hyphae transition can be induced by environmental factors, chemical signalling and metabolic adaptations, as well as cross-kingdom interactions with microbes[14, 15, 17–19]. Some of the bacterial species described elsewhere enhance the hyphae formation and adhesion of *C. albicans* to the oral cavity and bladder mucosa, increasing likely oral candidiasis and urinary tract infections[19]. Therefore, this transition is a crucial virulence trait for fungal pathogenesis but dispensable for growth and is considered a promising target for treating candidiasis by reducing pathogenesis[8].

Recent decades of research have highlighted probiotics play an important role against infectious pathogens through their effects on the epithelium, the production of antimicrobial compounds, and competitive exclusion[20–23]. Several efforts have been made to study the functional properties of cyclic dipeptides derived from microorganisms and to identify their effectiveness in human health as well as disease prevention [24, 25]. cyclo(L-Pro-D-Arg) identified from *Bacillus cereus* that showed antibacterial and antitumor activity[26]; cyclo(L-Leu-L-Tyr) derived from fungi *Penicillium* sp.[27] and cyclo(L-Leu-L-Pro) isolated from *Bacillus amyoliquefaciens* that inhibits *Streptococcus epidermidis* biofilm[28]. *Lactobacillus-*secreted compounds inhibited *C. albicans* growth and morphogenesis[10, 12]. Cyclo(L-Phe-L-Pro) and cyclo(L-Phe-trans-4-OH-L-Pro) produced by *Lactobacillus plantarum* (MiLAB 393) showed antifungal activity[29]. *Lactobacillus rhamnosus* produced molecule, 1-acetyl-β-carboline targets *C. albicans* YAK1 and inhibits yeast-to-hypha transition[10]. Administration of *Aeromonas veronii* as live or dietary feeds has protection against bacterial infections and improves host immunity as well as health[30, 31]. Gramicidin S is a cyclic decapeptide isolated from symbiotic *Aeromonas veronii* and showed antibacterial activity against *Dermacoccus nishinomiyaensis* and *Aeromonas hydrophila*[32]. Despite its antimicrobial activity, Gramicidin S is applied topically to treat superficial infections[33]. Recently, we reported that the culture supernatant of probiotic *A. veronii* V03 inhibited the growth of *Aeromonas hydrophila* and *Vibrio* species[34]. Moreover, dietary supplementation of *A. veronii* V03 enhanced the protection of *Cyprinus carpio* during *A. hydrophila* infection[30]. Nevertheless, the purification and structural characterization of cyclic dipeptides from greenish-yellow pigmented probiotic strain *A. veronii* V03 and their antimicrobial efficacy are largely unexplored. Hence, the current study aimed to isolate and identify structurally unique cyclic dipeptides from *A. veronii* V03 and to evaluate their antibacterial and antifungal activities using *in vitro* studies.

## 2.2. Materials and Methods

### 2.1. Bacterial strain and fermentation

Greenish-yellow pigmented probiotic strain *A. veronii* V03 was identified by 16S ribosomal RNA gene sequencing and showed desirable functional probiotic characteristics [34]. For isolation and structural characterization of bioactive compounds, strain V03 was grown in Luria-Bertani broth (LB, HiMedia, M1245) as described earlier[30, 34]. In brief, 50 ml overnight culture was transferred into 400 mL of sterile medium and incubated on a rotary shaker (120 rpm) at 30 °C for 72 h. Fermented cultures were centrifuged (6500×g, 15 min at –4 °C) and cell-free supernatants were obtained by filtration (Thermo Scientific, Waltham, MA) and stored at –4 °C until the extraction of metabolites.

### 2.2. Extraction and purification of bioactive compounds

Cell-free supernatants were extracted with an equal volume of ethylacetate (1:1 ratio) and the organic layer was condensed by a rotary evaporator (IKA-RV10, Switzerland)[35]. Dried residue yielded 2.5 g/15 L of culture filtrate. They were loaded onto silica gel using a glass column (30 cm × 2 cm) packed with silica (100-200 mesh; HiMedia). Separations were achieved by linear gradient elution using an initial composition of 98% chloroform and 2% methanol, then decreased stepwise as follows: chloroform: methanol (v/v) 95:5, 92:8, 85:15, 75:25, 70:30, 50:50, followed by 100% methanol was finally added to the column. Samples were dried and similar fractions were pooled according to their thin-layer chromatography (TLC, Silica gel 60 F_254_ plates, Merck) profile into six fractions (F1-F6) that included minor (F2, 265 mg; pooled F3-F4, 320 mg) and major (fraction 5, 631 mg, F6, 533 mg) factions were selected further purification based on their bioactivity. Secondary fractionation achieved by column chromatography (silica 200-400 mesh or C18) with linear gradient elution using hexane: ethyl acetate (v/v 80:20, 70:30, 50:50, 70:30) or chloroform/methanol similar conditions described above. ^1^H NMR spectra of both fractions yielded white crystalline compounds such as cyclo(L-Pro-L-Leu) (DKP-1), cyclo(L-Pro-L-Val) (DKP-2), cyclo(L-Pro-L-Phe) (DKP-3) and cyclo(L-Pro-L-Tyr) (DKP-4).

### 2.3. Structural characterization of bioactive compounds

Structural elucidation of isolated compounds was determined by nuclear magnetic resonance (NMR) spectroscopy using denudated chloroform with 0.1% tetramethylsilane as an internal standard. Proton (^1^H) and ^13^C NMR spectra of isolated compounds were acquired in a Bruker spectrometer (Bruker DRX-500 MHz, Rheinstetten, Germany) equipped with a cryogenically cooled prodigy broadband inverse detection probe (Bruker Biospin, Billerica, MA). Chemical shifts for CDCl_3_ δH 7.26 and δC 77.0 were expressed in parts per million (ppm) and the coupling constant *J* value was given in Hertz (Hz). Purified compounds were analyzed by high-resolution mass spectrophotometer (HRMS) and m/z values were acquired using the electrospray ionization mode (Orbitrap LC-MS, Thermo Scientific) equipped with XBridge BEH C18 column (Waters, Milford, MA). The specific optical rotation of isolated compounds was measured by a Jasco P-2000 digital polarimeter coupled with a sodium lamp (Na) at a wavelength of λ589 nm as we have described earlier[35].

### 2.4. Microbial strains and growth conditions

Microbial strains, *Bacillus cereus* (MTCC1305), *Bacillus subtilis* (MTCC2756), *Staphylococcus epidermidis* (MTCC435), *Staphylococcus simulans* (MTCC3610), *Mycobacterium smegmatis* (MTCC993), *Escherichia coli* (MTCC2622), *Klebsiella pneumoniae* (MTCC109), *Proteus mirabilis* (MTCC425), *Salmonella typhi* (MTCC3216), *Candida albicans* (MTCC277), *Candida tropicalis* (MTCC184), *Candida glabrata* (MTCC3019), and *Candida parapsilosis* (MTCC998) were obtained from the Microbial Type Culture Collection (MTCC, Chandigarh, India). *Staphylococcus aureus* (ATCC29213), *Aeromonas hydrophila* (ATCC7966) and *Pseudomonas aeruginosa* (ATCC27853) were purchased from the American Type Culture Collection Centre (ATCC, Rockville, MD, USA). Pathogenic bacteria and fungi strains were grown in Mueller–Hinton broth (MHB, HiMedia) or sabouraud dextrose broth (SDB, HiMedia) for 18 h at 30 °C, respectively. All microbial cultures were maintained in 50 % glycerol at -80 °C for further analysis.

### 2.5. Agar well diffusion assay

Antimicrobial activity of organic metabolic extracts including ethyl acetate, chloroform and hexane were tested against bacterial and fungal pathogens using agar well diffusion assay[36]. In brief, pathogenic bacteria and fungi strains were grown in 10 ml of MH and SD broth in similar conditions. Log phase (1×10^6^ cells/ml) pathogenic cultures were swabbed on the surfaces of respective agar plates (MHA and SDA) by sterile cotton swabs (HiMedia). Wells were then punched (6 mm) into the agar plates and 1mg/well of each metabolic extract was added into each well. DMSO-treated wells served as a vehicle control. Plates were incubated at 30 °C and observed for clear zones around the wells after 24 h.

### 2.6. Disk diffusion assay

Antimicrobial activity of identified four DKPs was tested against pathogenic bacteria (*P. mirabilis*, *S. aureus*, *A. hydrophila,* and *P. aeruginosa*) and fungi (*C. albicans*, *C. tropicalis*, *C. glabrata,* and *C. parapsilosis*) using disc diffusion assay[37, 38]. In brief, log-phase pathogenic cultures (1×10^6^ cells/ml) were swabbed on MHA and SDA plates. Vechile and DKPs (25 μg/disc dissolved in DMSO) treated disks were placed over them. DMSO treated disk served as vehicle control. After incubation for 24h, the growth of inhibition zones was measured in diameter.

### 2.7. Assessment of minimum inhibitory (MIC) and minimum bactericidal or fungicidal concentrations (MBCs or MFCs) of DKPs

MIC, MBC and MFC of DKPs were determined by the microdilution method[26, 37, 38]. Briefly, pathogenic bacteria (2×10^5^ cells/well) and fungi (2×10^5^ cells/well) were seeded into 96 well plates. They were treated with different doses (6.25 to 400µg/ml) of each DKP dissolved in culture media. Dimethyl sulfoxide (DMSO; 0.25%) treated cells served as a vehicle control. Plates were incubated at 30 °C for 24 h. Subsequently, 40 µl alamarBlue reagent (ThermoFisher Scientific, Waltham, MA) was added into each well and re-incubated for 1 h. Fluorescent intensity was measured according to manufacturer instructions. MIC, MBC and MFC were determined by the lowest concentrations inhibiting 50% growth or no growth in tested bacterial or fungal pathogens, respectively.

### 2.8. Effects of DKPs on C. albicans yeast-hyphae transition

The inhibitory effects of DKPs on yeast-hyphae formation of *C. albicans* were determined by using the previously described method[39]. Loopful colonies from freshly streaked *C. albicans* on SDA plate were inoculated into 10 ml of NYP media (containing N-acetylglucosamine 10 mol/l, yeast nitrogen base 3.35 g/l, proline 10 mol/l, NaCl 4.5 g/l, and pH 5.4 ± 0.2). Briefly, 2×10^6^ cells/ml were treated without or with isolated DKPs (12.5 to 150 μg/ml) and incubated for 3 h at 30 °C. Cells were washed with PBS and fixed with 4% paraformaldehyde. Yeast-to-hyphae transition of *C. albicans* was observed and images were acquired using the light microscope (Olympus Binocular Microscope-CX21, under 20X magnification) equipped with a digital camera HDCE-X3 supported with professional imaging software (Scope image-9.0). Yeast-to-hyphae length was quantified using Image-J software.

### 2.9. Effects of DKPs on phospholipase (PL) and secreted aspartyl proteinase (SAP) activity

Effects of DKPs on PL and SAP activity of *C. albicans* were evaluated using the previously described method[40–42], with minor modifications. Concisely, 2×10^6^ cells/ml were treated without or with isolated DKPs (12.5 to 150 μg/ml) and incubated at 30 °C for 1h. Treated cells were washed and re-suspended in PL (containing (g/l) NaCl 57.3, peptone 10, glucose 30, CaCl_2_ 0.55, and sterile egg yolk emulsion 8%) and SAP inducing media (containing (g/l) yeast carbon base 23.4, yeast extract 2, bovine serum albumin 4, and pH 5.0). They were incubated at 30 °C for 72 h followed by the PL and SAP activity was determined using the method described previously[42, 43]. PL and SAP activity was calculated by the following formula: Percentage of activity (%) = [(test/control) ×100)].

### 2.10. Determination of DKPs on C. albicans biofilm formation

Biofilm formation of *C. albicans* in response to DKPs treatments was analyzed using the previously reported method[11, 44]. In brief, 5×10^5^ cells/well were treated without or with isolated DKPs (12.5 to 150 μg/ml) in RPMI-1640 supplemented with MOPS. The culture plate was incubated for 24h at 37°C to permit biofilm formation. The inoculum was removed and biofilms were washed three times with PBS (to remove planktonic/unattached cells). Biofilm formation was fixed and quantified using the crystal violet reporter assay[10, 11].

### 2.11. Statistical analysis

All statistical analysis was performed using GraphPad Prism statistical software (San Diego, CA, USA). Data were analyzed by one-way analysis of variance (ANOVA) with Dunnett’s multiple comparison test and *p <* 0.05 was considered statistically significant. All data were presented as mean ± standard deviation (SD) of four biological replicates.

## 3. Results

### 3.1. Purification of bioactive metabolites

We reported *A. veronii* V03 culture supernatants exhibited promising antibacterial activity[34]. Therefore, we employed a bioactivity-guided purification strategy to identify which metabolite(s) were responsible for observed inhibitory activity. Greenish-yellow extracts, including ethyl acetate, chloroform, and hexane were obtained from the cell-free culture filtrate of *A. veronii* V03. They were tested for their possibility of antimicrobial activity against Gram-positive and Gram-negative bacterial and fungal pathogens by agar well diffusion assay. Out of 3 metabolic extracts, only ethyl acetate extract-derived metabolites exhibited broad-spectrum antimicrobial activity against tested bacterial and fungal pathogens (Table S1 and S2). Particularly, ethyl acetate-derived metabolites showed predominant activity against Gram-negative bacteria compared with Gram-positive and fungal pathogens (Table S1 and S2). Out of 14 bacterial pathogens, only 4 bacterial pathogens exhibited the highest zone of inhibition, 24.5 mm for *P. mirabilis*, 24 mm for *S. aureus,* 22.5 mm for *A. hydrophila*, and 22 mm for *P. aeruginosa*, respectively (Table S1). The ethyl acetate extract was fractioned and tested for its bioactivity against *P. aeruginosa* and *C. albicans*. Out of six fractions, fractions F2-6 exhibited promising antimicrobial activity against tested pathogens (Table S3; Fig. S1A, B). Separated bioactive fractions yielded four cyclic dipeptides eluted at 35% ethyl acetate in hexane (DKP-1), 50% ethyl acetate in hexane (DKP-2), 2% methanol in chloroform in hexane (DKP-3), and 15% methanol in chloroform (DKP-4). TLC revealed that eluted cyclic dipeptides were single spots.

### 3.2. Structural elucidation of bioactive compounds

Structures of purified DKPs were elucidated by ^1^H and ^13^C NMR and mass spectroscopy techniques. Cyclic dipeptides were isolated and identified as cyclo(L-Pro-L-Leu) (DKP-1), cyclo(L-Pro-L-Val) (DKP-2), cyclo(D-Pro-L-Phe) (DKP-3), and cyclo(L-Pro-D-Tyr) (DKP-4) (Fig. 1). As shown in ^1^H and ^13^C NMR spectrum of DKPs were showed triplet signals at δH 4.01 ppm and δC 59.0 ppm, along with multiplets at δH 2.2, 1.9, and 3.4 ppm represents the presence of proline (Fig. S2). Amide hydrogen signal appeared at 5.9 ppm in the NMR spectra of isolated DKPs that had proline residue. The absolute configuration of purified DKPs was determined by comparing our results with previously reported specific optical rotation values in the literature[35, 45–47].

**Figure 1.**
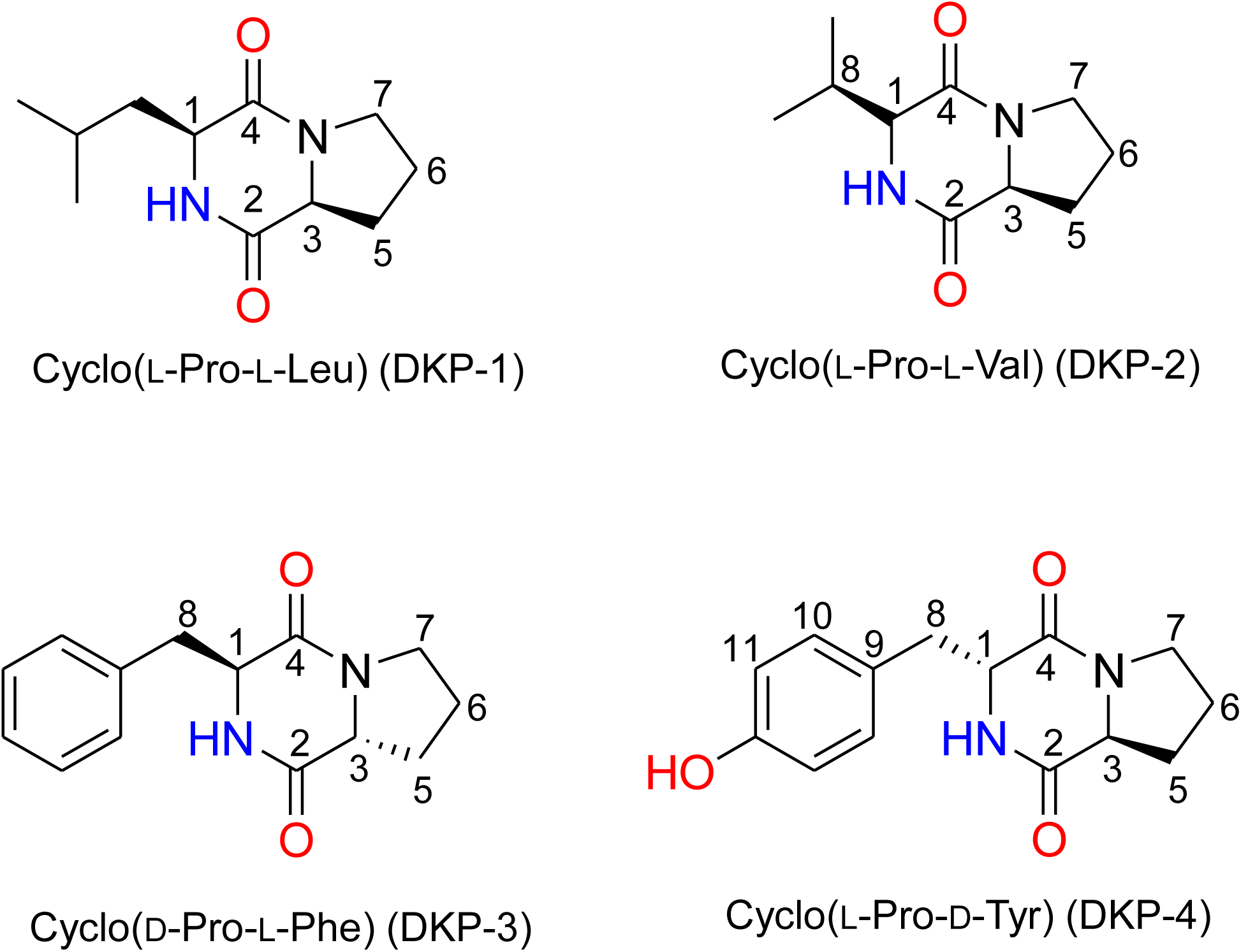
Structures of cyclic dipeptides from *Aeromonas veronii* V03. Chemical structures of four cyclic dipeptides were identified from *E. acetylicum* S01.

DKP 1: Cyclo (L-Pro-L-Leu): White crystalline powder 33 mg; TLC (methanol: chloroform, 1:99 v/v): Rf *=* 0.9; HPLC (methanol (MeOH): acetonitrile (CH_3_CN), 70:30 v/v): RT = 0.91 min; Specific Optical Rotation: [α]^26^_D_ - 109° (*c* = 0.5, ethanol), for R,R [α]^24^_D_ = - 109° (*c* = 0.4, ethanol)[45]; ^1^H and ^13^C NMR data, see Table 1 (Fig. S2A, B); HR-MS (m/z): [M + H]^+^ calcd for C_11_H_19_N_2_O_2_, 211.1450; found, 211.1448 (Fig. S3A).

**Table 1:**
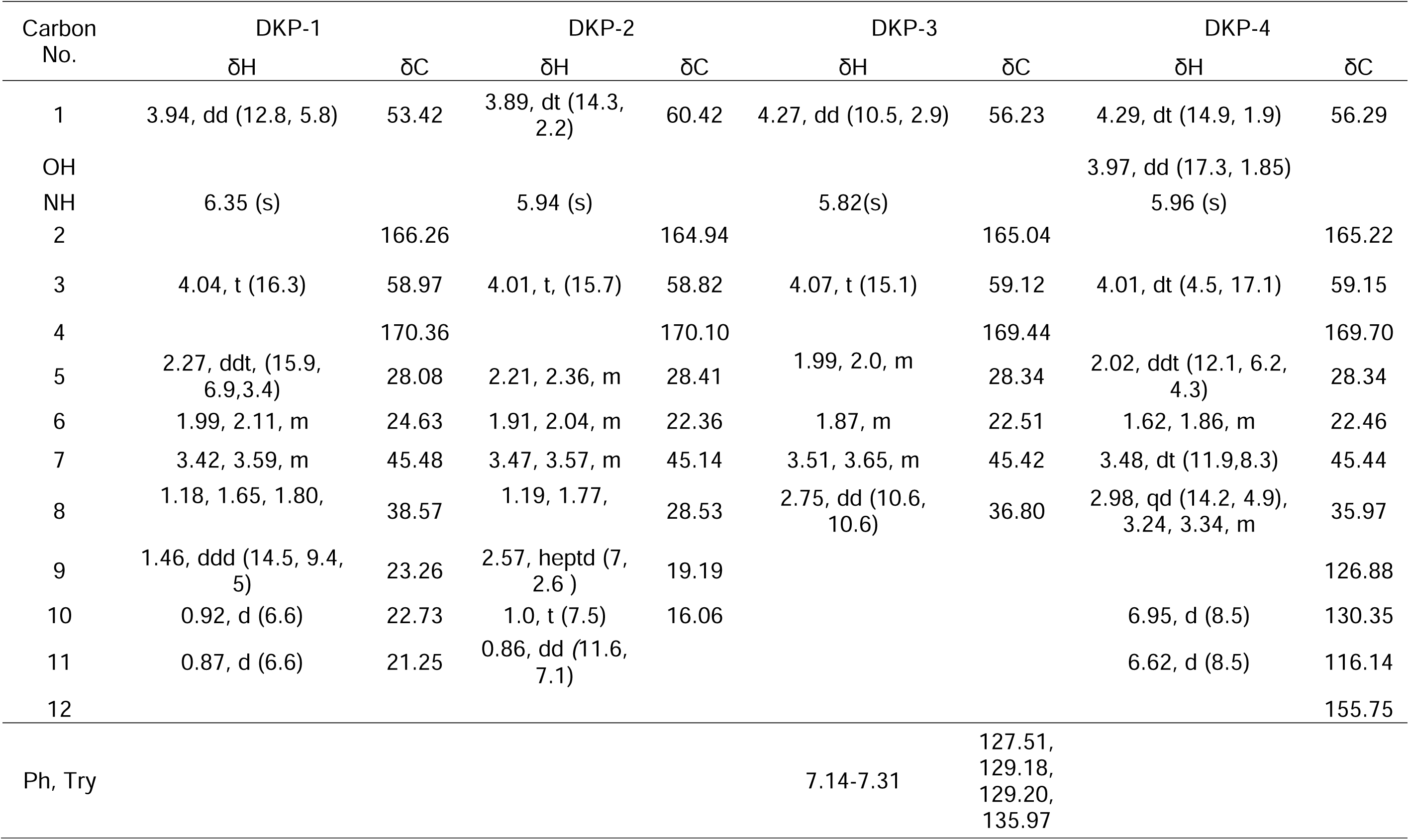
NMR data for DKPs 1 – 4 in CDCl_3_ (multi, *J,* Hz)

DKP 2: Cyclo (L-Pro-L-Val): Whilte powder 35.2 mg; TLC (ethyl acetate: hexane, 60:40 v/v): Rf = 0.3; HPLC (MeOH: CH_3_CN, 70:30 v/v): RT = 0.89 min; Specific Optical Rotation: [α]^24^_D_ = -137° (*c* = 0.2, ethanol), for R,R, [α]^24^_D_ -139.4° (*c* = 0.16, EtOH)[46]; ^1^H and ^13^C NMR data, see Table 1 (Fig. S2C, D); HR MS (m/z): [M + H]^+^ calcd for C_10_H_17_N_2_O_2,_ 197.1292; found, 197.1292 (Fig. S3B).

DKP 3: Cyclo (D-Pro-L-Phe): Pale yellow powder (23 mg); TLC (ethyl acetate: hexane, 85:15 v/v): Rf = 0.45; HPLC (MeOH: CH_3_CN, 60:40 v/v): RT value 0.94 min; Specific Optical Rotation: [α]^24^_D_ = +87° (*c* =0.2, MeOH), for R,R, [α] ^30^_D_ = +70.2° (*c* = 1.45, MeoH)[48]; ^1^H and ^13^C NMR data, see Table 1 (Fig. S2E, F); HRMS (m/z): [M + H]^+^ calcd for C_14_H_17_N_2_O_3,_ 245.1292; found, 245.1291 (Fig. S3C).

DKP 4: Cyclo (L-Pro-D-Tyr): Pale yellow powder (38.3 mg); TLC (methanol: chloroform, 1:99 v/v): Rf = 0.80; HPLC (MeOH:CH_3_CN, 50:50 v/v): RT = 0.99 min; Specific Optical Rotation: α]^24^_D_ = -113° (*c* = 0.2, EtOH), for R,R,[α]^30^_D_ = -118°(c = 0.11, EtOH)[47, 48]; ^1^H and ^13^C NMR data, see Table 1 (Fig. S2G, H); HRMS (m/z): [M + H]^+^ calcd for C_14_H_17_N_2_O_3_, 261.1241; found, 261.1243 (Fig. S3D).

### 3.3. Antimicrobial activity of DKPs against fungal and bacterial pathogens

Disk diffusion assay revealed that identified DKPs showed promising antimicrobial activity against tested bacterial and fungal pathogens (Table 2). It was found that *P. mirabilis* was more susceptible to all DKPs at a dose of 25 μg/ml compared to other bacterial pathogens (Table 2). Additionally, the DKP-3 exhibited the highest zone of inhibition against tested fungal pathogens compared to other DKPs (Table 2).

**Table 2:** Antibacterial and antifungal activities of cyclic dipeptides (DKPs) identified from *A. veronii* V03.

| Microorganisms | DKP-1 | DKP-2 | DKP-3 | DKP-4 |
| --- | --- | --- | --- | --- |
| Zone of inhibition (mm with Disc) [25µg/Disc] |  |  |  |  |
| Bacterial pathogens |  |  |  |  |
| *P. mirabilis* | 23 ± 1 | 24 ± 0 | 26.5 ± 0.5 | 23 ± 0 |
| *A. hydrophila* | 11 ± 0 | 14.5 ± 0.5 | - | - |
| *P. aeruginosa* | - | 11 ± 1 | 10 ± 0 | - |
| *S. aureus* | ++ | ++ | 8 ± 0 | 11 ± 1 |
| Fungal pathogens |  |  |  |  |
| *C. albicans* | 15±0.5 | 10 ± 0 | 18 ± 0 | 20 ± 1 |
| *C. tropicalis* | ++ | 9.5 ± 0.5 | 16.5±0.5 | 13±0 |
| *C. glabrata* | ++ | 11 ± 1 | 14 ± 0.5 | 11 ± 0 |
| *C. parapsilosis* | 8 ± 0 | ++ | 17 ± 0 | 10.5 ± 0.5 |

Results are shown as mean ± SD; DKP-1: cyclo(L-Pro-L-Leu), DKP-2: cyclo(L-Pro- L Val), DKP-3: cyclo(D-Pro- L-Phe); DKP-4: cyclo(L-Pro- D-Tyr). – = No inhibition, +/++ = static inhibition; mm – millimeter

### 3.4. Minimum inhibitory and minimum bactericidal or fungicidal concentrations of DKPs

MIC and MBC of DKPs against tested bacterial pathogens (Table 3). We noted the lowest MIC (7.85 μg/ml) and MBC (19.41 μg/ml) of DKP-3 against *P. mirabilis* compared to other DKPs. DKP-1 exhibited the lowest MIC (20.62 μg/ml) and MBC (87.5 μg/ml) against *A. hydrophila* compared to other DKPs. We found promising inhibitory activity of DKP-3 and DKP-4 against *S. aureus* and DKP-3 against *P. aeruginosa.* All DKPs showed more effectiveness against *P. mirabilis* compared to other bacterial pathogens. The MIC and MFC of isolated four DKPs against fungal pathogens (Table 4). Consistent with antibacterial activity, DKP-3 demonstrated promising antifungal activity with the lowest MIC at 10.62 to 15.88 µg/ml against tested fungal pathogens (Table 4).

**Table 3.** The MICs/MBCs of DKPs were determined against bacterial pathogens.

| Bacterial pathogens | MIC/MBC (µg/ml) |  |  |  |  |  |  |  |
| --- | --- | --- | --- | --- | --- | --- | --- | --- |
|  | DKP-1 |  | DKP-2 |  | DKP-3 |  | DKP-4 |  |
|  | MIC | MBC | MIC | MBC | MIC | MBC | MIC | MBC |
| *P. mirabilis* | 10.62 | 25.21 | 10.52 | 19.87 | 7.85 | 19.41 | 11.5 | 27.91 |
| *A. hydrophila* | 20.62 | 87.5 | 21.87 | 93.25 | 199.3 | >200 | 119 | 137.5 |
| *P. aeruginosa* | 187.9 | >200 | 79.25 | 125.25 | 90.25 | 152.5 | 190.5 | >300 |
| *S. aureus* | 191.5 | >200 | 170.2 | >200 | 59.51 | 99.9 | 40.87 | 87.5 |

DKP-1: cyclo(L-Pro-L-Leu); DKP-2: cyclo(L-Pro- L-Val); DKP-3: cyclo(D-Pro- L-Phe); DKP-4: cyclo(L-Pro- D-Tyr). >200: resistance up to 350 µg/ml

**Table 4.** The MICs/MFCs of DKPs were determined against fungal pathogens.

| Fungal pathogens | MIC/MFC (µg/ml) |  |  |  |  |  |  |  |
| --- | --- | --- | --- | --- | --- | --- | --- | --- |
|  | DKP-1 |  | DKP-2 |  | DKP-3 |  | DKP-4 |  |
|  | MIC | MFC | MIC | MFC | MIC | MFC | MIC | MFC |
| *C. albicans* | 31.62 | 69.59 | 48.87 | 105.62 | 10.62 | 33.89 | 51.25 | 100.59 |
| *C. tropicalis* | 21.71 | 45.21 | 87.91 | 124.55 | 15.89 | 38.5 | 95.21 | 129.1 |
| *C. glabrata* | 62.26 | 110.52 | 57.55 | 99.62 | 19.62 | 43.8 | 80.69 | 102.32 |
| *C. parapsilosis* | 72.5 | 137.65 | 47.87 | 81.25 | 15.88 | 37.52 | 51.6 | 98.81 |

DKP-1: cyclo (L-Pro-L-Leu); DKP-2: cyclo (L-Pro- L-Val); DKP-3: cyclo (D-Pro- L-Phe); DKP-4: cyclo (L-Pro- D-Tyr)

### 3.5. DKPs inhibit yeast-hyphae formation

The morphological transition is important for *C. albicans* to infect humans and cause disease[16]. Thus, the effects of DKPs on the *C. albicans* yeast-to-hyphae transition were evaluated under hyphal induction conditions. As evident from the images, the majority of *C. albicans* cells in the untreated group had formed germ tubes, while hyphal formation was inhibited in a dose-dependent manner by exposure to DKPs (Fig. 2A-B). All DKPs reduced hyphal formation in *C. albicans* by more than 70 % at a concentration of 50 µg/ml. While *C. albicans* treated with higher concentrations of DKPs showed 100 % inhibition of hyphal formation and cells underwent shrinking, membrane rupture, and death (Fig. 2A-B).

**Figure 2.**
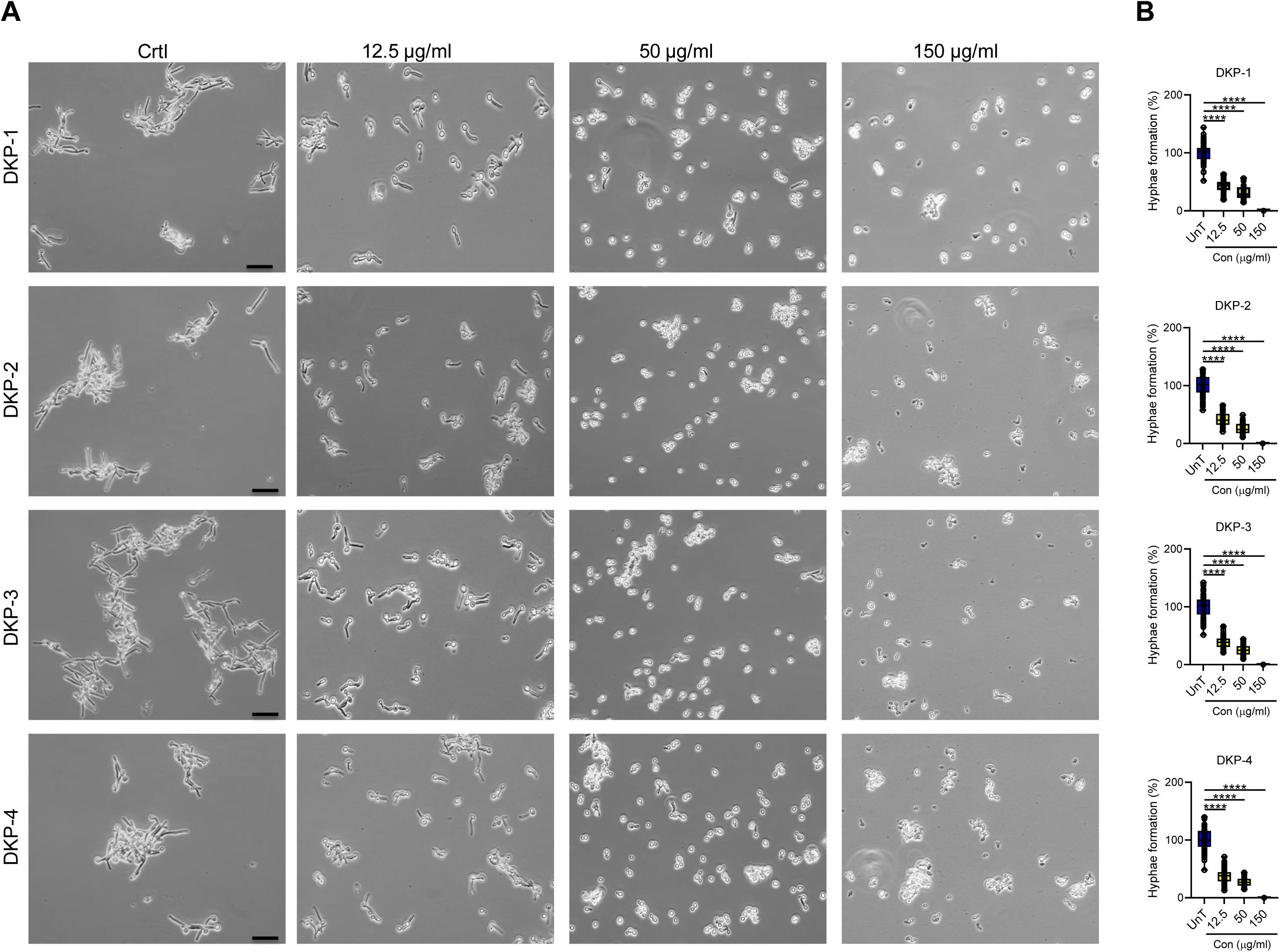
Cyclic dipeptides inhibits *Candida albicans* yeast-hypha transition. **A.** Representative pictures of yeast-hyphae transition of untreated and treated with DKPs for 3h. **B**. Quantification of the hyphae length of *C. albicans*. The percentage of hyphae formation was determined relative to the untreated (UnT) cells. Data were expressed as mean ± SD (n = 4). One-way ANOVA with Dunnett’s multiple comparison test. The asterisks \**p* < 0.05, \*\**p* < 0.001, \*\*\**p* < 0.0001, \*\*\*\**p* < 0.0001, indicate a significant difference between the UnT in response to DKPs treated cells. Images were acquired using an Olympus Binocular Microscope – CX21, under 20X magnification). DKP 1 – cyclo(L-Pro-L-Leu), DKP 2– cyclo(L-Pro-L-Val), DKP 3 – cyclo(D-Pro-L-Phe), and DKP 4 – cyclo(L-Pro-D-Tyr).

### 3.6. Effects of DKPs on PLs and SAPs enzymatic activity of C. albicans

To evaluate the effects of DKPs on the pathogenicity of *C. albicans*, we then tested whether these compounds influenced *C. albicans* secreted hydroxylase. Tested four DKPs inhibited PLs and SAPs of *C. albicans* in a dose-dependent manner compared to untreated cells (Fig. 3A-B). The percentage of PL activity was reduced by 75.3% for DKP-3 at a concentration of 150 µg/ml (Fig. 3A; *p* = 0.0001). Consistent with PLs activity, SAPs were significantly reduced by at least 60% in DKPs-treated *C. albicans* relative to untreated cells (*p* = 0.00001; Fig. 3B).

**Figure 3.**
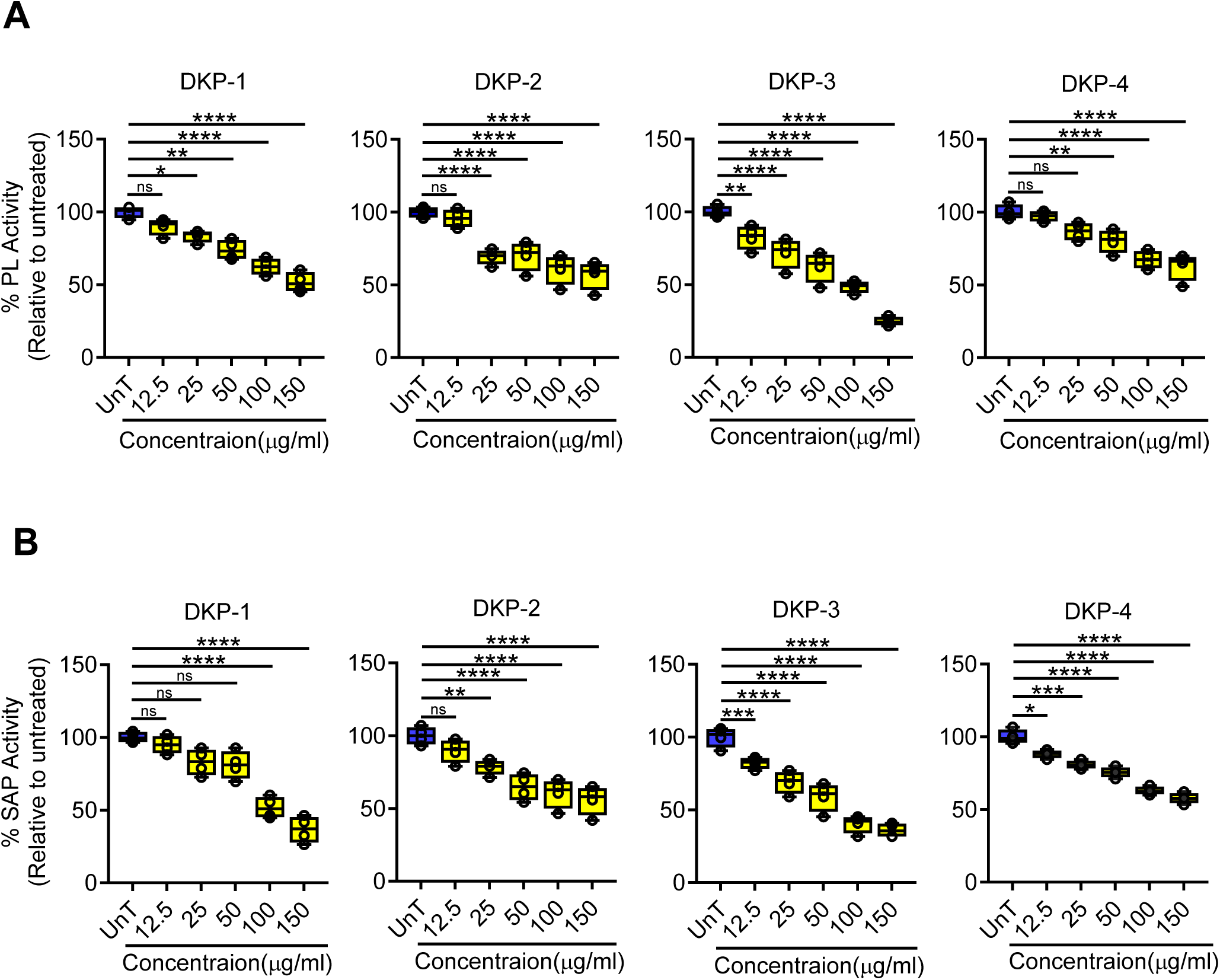
Cyclic dipeptides inhibits secreted hydrolases of *Candida albicans*. **A**. Quantification of phospholipase (PLs) activity. **B.** Quantification of secreted aspartic protease (SAPs) activity. The percentage of PLs and SAPs was determined relative to the untreated (UnT) cells. Data were expressed as mean ± SD (n = 4). One-way ANOVA with Dunnett’s multiple comparison test. The asterisks \**p* < 0.05, \*\**p* < 0.001, \*\*\**p* < 0.0001, \*\*\*\**p* < 0.0001, indicate a significant difference between the UnT in response to DKPs treated cells. DKP 1 – cyclo(L-Pro-L-Leu), DKP 2– cyclo(L-Pro-L-Val), DKP 3 – cyclo(D-Pro-L-Phe), and DKP 4 – cyclo(L-Pro-D-Tyr).

### 3.7. DKPs inhibit C. albicans biofilm formation

The formation of biofilms by *C. albicans* on the mucosa or endothelium in humans or on the surface of implemented medical devices has been linked to recurrent infections and treatment failure with antifungal therapy[17]. We, therefore, tested the effects of DKPs to inhibit the formation of biofilms by *C. albicans* using the standard method described previously[10, 11]. All DKPs exhibited a dose-dependent inhibition of *C. albicans* biofilm formation (Fig. 4A-B). Against *C. albicans,* at 150µg/ml DKP-3 and DKP-1 increased biofilm inhibition by 79% and 83%, relative to the untreated cells (Fig. 4B).

**Figure 4.**
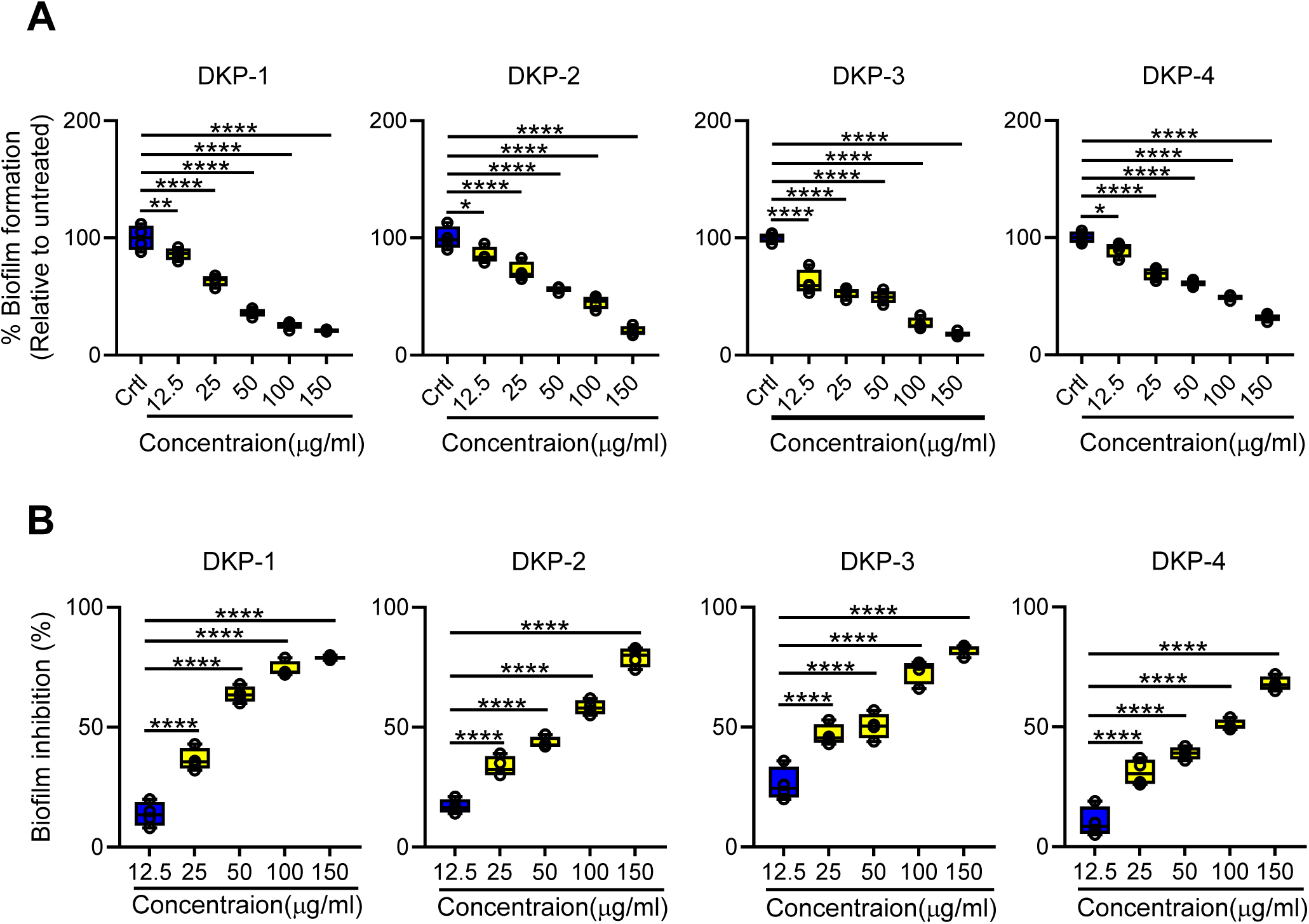
Cyclic dipeptides inhibits *Candida albicans* biofilm formation. **A.** Percentage of the biofilm formation. **B.** Inhibition of *C. albicans* biofilm formation. Inhibition of biofilm formation by cyclic dipeptides was quantitatively assessed using a crystal violet assay. The percentage of biofilm inhibition was determined relative to the untreated (UnT) cells. Data were expressed as mean ± SD (n = 4). One-way ANOVA with Dunnett’s multiple comparison test. The asterisks \**p* < 0.05, \*\**p* < 0.001, \*\*\**p* < 0.0001, \*\*\*\**p* < 0.0001, indicate a significant difference between the UnT in response to DKPs treated cells. DKP 1 – cyclo(L-Pro-L-Leu), DKP 2– cyclo(L-Pro-L-Val), DKP 3 – cyclo(D-Pro-L-Phe), and DKP 4 – cyclo(L-Pro-D-Tyr).

## 4. Discussion

Bacterial and fungal infections remain a significant concern for human health[1, 2]. Increasing drug resistance among pathogens leads to extended hospitalization, rising treatment costs and increased morbidity and mortality[1, 7, 8]. Thus, new therapeutic approaches are required to address the global challenge of drug-resistant microbial infections[2, 8, 9, 23]. Several studies have reported that probiotics are used to treat microbial infections, mostly bacterial and fungal pathogens[12, 20, 21]. The fundamental truth is that probiotic–derived metabolites, including antimicrobial peptides and organic acids, play a critical role in the prevention of infectious pathogens via their antimicrobial effects on host mucosal surfaces[49]. Notably, cyclic dipeptides are widely produced by microorganisms and are important components of host immunity[24, 25, 49]. These inherent functions have health-promoting and disease-preventing functions[10, 49]. Cyclic decapeptide produced by *A. veronii* showed antibacterial activity against human pathogens, *D. nishinomiyaensis* and *A. hydrophila*, but not against *A. vernoii* itself[32]. Recent studies also showed the supreme activity of gramicidin S against staphylococci and enterococci pathogens that inhibits its biofilm[50]. Nevertheless, only limited information is available on the chemical characterization of antimicrobial peptides from *Aeromonas* spp.

In this study, four cyclic dipeptides (DKP 1-4) were isolated and identified from *A. veronii* V03. We found that four DKPs exhibited promising broad-spectrum antimicrobial activity against bacterial and fungal pathogens. Importantly, *P. mirabilis* was more susceptible to DKP treatments compared to other pathogens. *P. mirabilis* an opportunistic pathogen that causes various urinary tract infections, such as cystitis, pyelonephritis, and bacteriuria, especially in patients with diabetes. It also causes bacteriemia and progresses to life-threatening urosepsis[3]. Besides, DKP-3 exhibited potent inhibitory activity against *C. albicans* but also on non-albicans species. Subtle structural variances are known to play a crucial role in the inhibition of these four peptides. Our finding was consistent with previous studies, cyclo(L-Pro-L-Phe) and cyclo(L-Phe-trans-4-OH-L-Pro) from *L. plantarum* MiLAB 393 that showed antimicrobial activity against bacterial and fungal pathogens[29]; cyclo(L-Pro-D-Arg) isolated from *B. cereus* inhibited *P. mirabilis* growth[26]; cyclo(L-Pro-L-Leu) and cyclo(L-Pro-L-Tyr) that inhibited *C. albicans* growth[48]. We also observed four DKPs showed a substantial effect against *P. aeruginosa* and *S. aureus*. Li and co-workers reported that cyclo(L-Tyr-L-Pro) and cyclo(L-Tyr-D-Pro) derived from *Lactobacillus reuteri* while cyclo(L-Tyr-L-Pro) exhibited the highest activity against *S. aureus*[51]. Another study also reported that cyclo(L-Tyr-L-Pro) and cyclo(L-Pro-L-Phe) inhibited biofilm formation and quorum sensing signalling of *P. aeruginosa* PA01[4, 52]. Smaoui and co-workers reported the antimicrobial activity of cyclo(L-Pro-L-Tyr) against both Gram-positive and Gram-negative bacteria besides fungus[53]. We and others reported that DKP-3 demonstrate potent antifungal activity against *C. albicans*, which is better than the standard fungicide Amphotericin B[54]. The possible explanation for low and high peptide concentrations required for significant growth inhibition of microbial pathogens described above is due to several factors including cell wall modifications, efflux pumps or lack of sufficient assimilation of peptides into the cells[55, 56].

Fungal cell walls are dynamic structures that are essential for cell viability, morphogenesis, and pathogenesis. Also, *C. albicans* has various virulence factors such as the morphological transition from yeast to hyphae, secretion of hydrolytic enzymes, the expression of adhesions and invasions on the cell surfaces, and biofilm formation[14, 15]. Particularly, various hydrolytic enzymes, such as SAPs and LPs secreted by *C. albicans* hyphae contribute to the tissue invasion process[16, 18, 40]. Current antifungals interfere with cellular processes that are essential for fungal growth and thus easily stimulate the development of drug resistance by *C. albicans*[8, 16, 56]. However, antivirulence agents are less likely to result in drug resistance and perturb the healthy microbiota[7, 8, 10]. Therefore, targeting the virulence traits of *C. albicans* is a promising therapeutic strategy to reduce pathogenesis and overcome antifungal resistance[9]. Hitherto, the antivirulence effects of DKPs against *C. albicans* remain unknown. Here, we report that DKPs produced by *A. veronii* V03 inhibited important virulence traits of *C. albicans,* yeast-to-hypha transition and secreted hydrolases, SAPs and PLs. These results suggest DKPs can attenuate *C. albicans* pathogenesis by inhibiting hyphae formation and secreted hydrolases. It has been reported by the National Institutes of Health that biofilms contribute to approximately 80% of all microbial infections in the United States[16, 37]. Current approaches to inhibit *C. albicans* biofilm formation in healthcare settings such as catheters include coating the surface with antifungal agents whereby high concentrations of an antifungal agent slowly diffuse via the lumen of a catheter before insertion of the catheter into a patient[2, 14, 17, 19]. Here, we show that DKPs were more effective at inhibiting *C. albicans* biofilm formation. Our results were well correlated with the previous studies, cecropin-4-derived peptide suppressed virulence of *C. albicans*, which includes hyphal transition and biofilm formation [57]. *Lactobacillus*-derived 1-acetyl-β-carboline prevents yeast-to-hyphae growth and biofilm formation of *C. albicans* via inhibition of Yak1[10]. Moreover, the inhibitory effect of tyrocidines on *C. albicans* biofilm formation is attributed to their disruptive impact on membrane integrity [58]. Antifungal peptide metchnikowin inhibits cell wall synthesis and integrity of *C. albicans* by inhibiting β(1,3)-glucanosyltransferase[59]. In conclusion, we report four DKPs were isolated and identified from *A. veronii* V03. This study revealed that four DKPs remarkably inhibited both bacterial and fungal pathogens. Besides, four DKPs inhibited virulence traits and biofilm formation of *C. albicans*. Collectively, our results suggest that DKPs would be promising antivirulence agents and support the notion that the antivirulence approach should be an integral part of developing therapies for the prevention and treatment of infectious caused by *C. albicans* or other *Candida* species that undergo yeat-hyphal transition. Further studies are required to explore underlying molecular mechanisms of action on DKPs against *C. albicans* and *P. mirabilis*.

## Acknowledgements

The authors wish to acknowledge the DST-Science Engineering Research Board, DST-SERB (Ref. SB/YS/LS-05/2014), Government of India, for providing financial support and DST-PURSE Phase-II and UGC-SAP for equipment support. The authors would like to thank Dr. L. Ravi Shankar and Mr. T. Arun Kumar from the Chemical Science and Technology Division, CSIR-National Institute for Interdisciplinary Science and Technology, Thiruvananthapuram, India, for permitting Dr. JS to use their laboratory facility for purification and characterization cyclic dipeptides. The authors thank the Ministry of Science and Technology of Taiwan and Kaohsiung Medical University, for providing a fellowship to Dr. SJ.

