## Supplementary File for "Inhibition of *Candida albicans* virulence factor by cyclic dipeptides derived from *Aeromonas veronii* V03"

**_Table S1._** _Antimicrobial activity of organic extract-derived metabolites from_ *_A. veronii_* _V03 against bacterial pathogens_

| **Test microorganisms** | **Zone of inhibition (1mg/well) includes well diameter (mm)** | | |
| --- | --- | --- | --- |
|  | **Ethyl acetate** | **Chloroform** | **Hexane** |
| *B. cereus^a^* | 14.5 ± 0.5 | 9.0 ± 0 | - |
| *E. coli^a^* | 12.5 ± 0.5 | - | - |
| *K. pneumoniae^a^* | 18.5 ± 0.5 | 11.5 ± 0.5 | - |
| *M. smegmatis^a^* | 13 ± 0 | ++ | - |
| *P. mirabilis^a^* | 20.5 ± 0.5 | 15 ± 0 | 12.5± 0.5 |
| *S. aureus^a^* | 13 ± 0 | - | - |
| *S. epidermidis^a^* | 11.5 ± 0.5 | - | 9 ± 0 |
| *S. simulans^a^* | 13 ± 0 | - | - |
| *A. hydrophila^b^* | 15 ± 0 | 10.5 ± 0.5 | - |
| *P. aeruginosa*^b^ | 14.5 ± 0.5 | ++ | - |
| *S. aureus^b^* | 14 ± 0 | - | - |
| *M. smegmatis^c^* | 13 ± 0 | - | 10.5± 0.5 |

Results are shown as mean ± SD, mm – millimetre, - No activity up to 2mg/well, ++ Static inhibition. ^a^MTCC culture, ^b^ATCC strains, ^c^NIRT strains.

**Table S2.** Antifungal activity of different organic extract–derived metabolites from *A. veronii* V03 against *Candida* sp.

| **Test microorganisms** | **Zone of inhibition (1mg/well) includes well diameter (mm)** | | |
| --- | --- | --- | --- |
|  | **Ethyl acetate** | **Chloroform** | **Hexane** |
| *C. albicans* MTCC277 | 13 ± 0 | ++ | ++ |
| *C. tropicalis* MTCC184 | 10.5 ± 0.5 | - | - |
| *C. glabrata* MTCC3019 | 10.5 ± 0.5 | - | - |
| *C. parapsilosis* MTCC998 | 12.5 ± 0.5 | - | 9.5 ± 0.5 |

Results are shown as mean ± SD, mm – millimetre, - No activity up to 2mg/well, ++ Static inhibition

**_Table S3._** _Antibacterial activity of partially purified fractions (F1-F6) from_ *_A. veronii_* _V03 against bacterial pathogens._

| **Test Pathogens** | **Zone of inhibition (250 µg/well) includes well diameter (mm)** | | | | | |
| --- | --- | --- | --- | --- | --- | --- |
|  | **F1** | **F2** | **F3** | **F4** | **F5** | **F6** |
| ***P. aeruginosa*** MTCC | - | 10.5 ± 0.5 | 11 ± 0 | 23 ± 0 | 25 ± 0 | 25± 0 |
| ***C. albicans ATCC*** | - | 12.5 ± 0.5 | 12± 1 | 10.5± 0.5 | 15± 0.5 | 16± 1 |

Results are shown as mean ± SD, mm – millimetre, - No activity, ++ Static inhibition, F –Fraction,

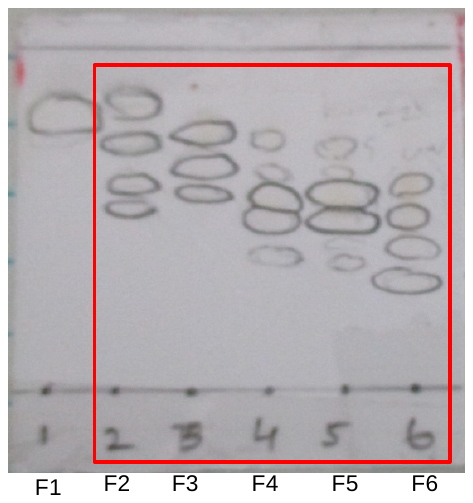

**Figure 1A**: TLC profile shows fractions (F1-F6)

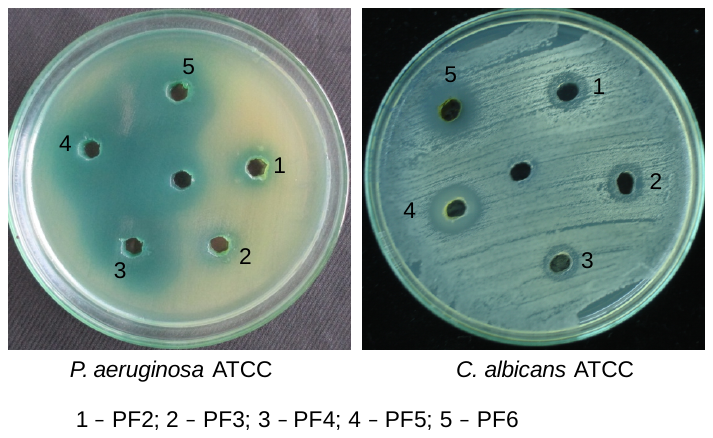

**Figure 1B**: Antimicrobial activity of partially purified fractions (F2-F6) against bacterial pathogens using agar well diffusion assay.

**NMR spectra of** **identified compounds from** *A. veronii* V03

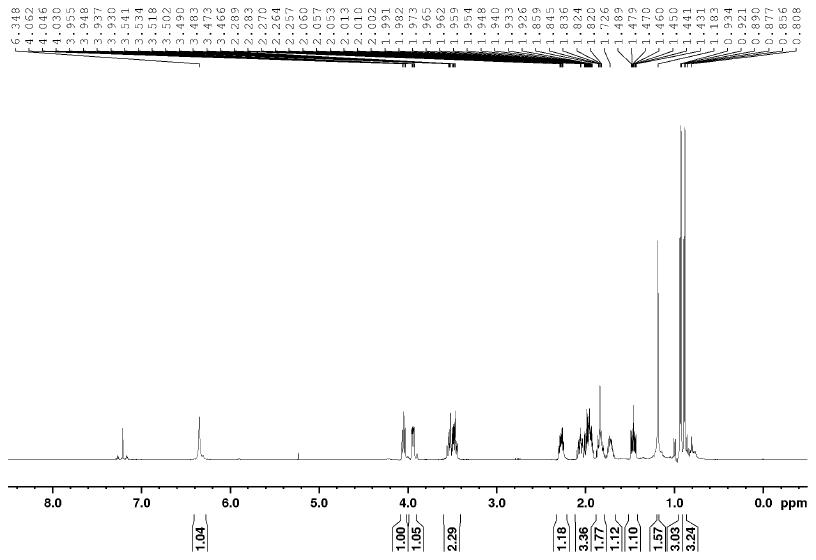

**Figure S2A. ^1^H NMR spectrum of DKP-1 (500 MHz; CDCl_3_)**

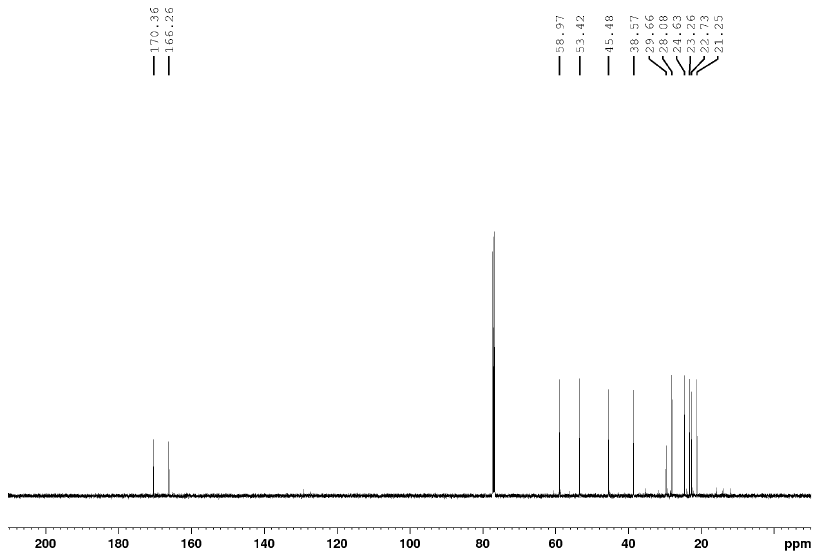

**Figure S2B. ^1^C NMR spectrum of DKP-1 (500 MHz; CDCl_3_)**

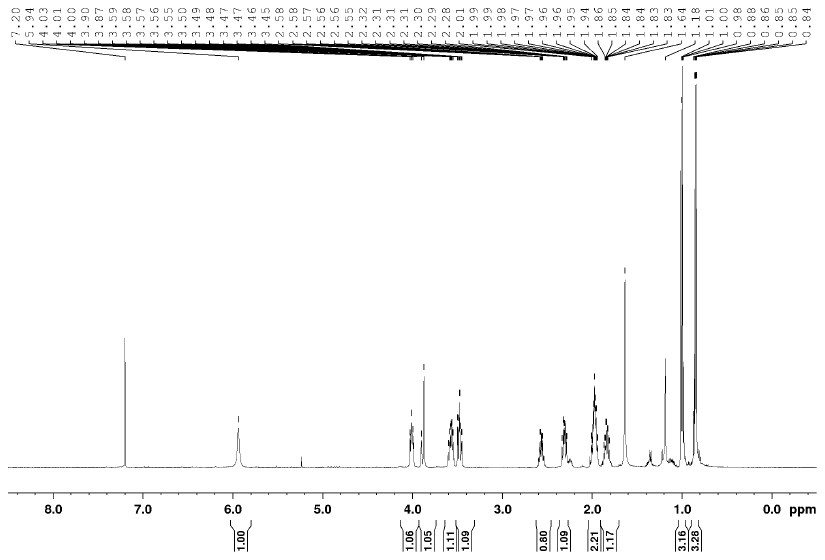

**Figure S2C. ^1^H NMR spectrum of DKP-2 (500 MHz; CDCl_3_)**

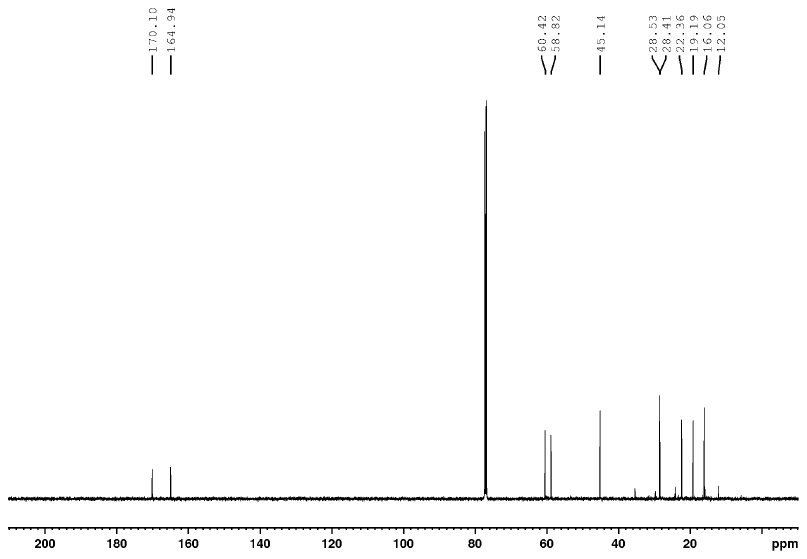

**Figure S2D. ^1^C NMR spectrum of DKP-2 (500 MHz; CDCl_3_)**

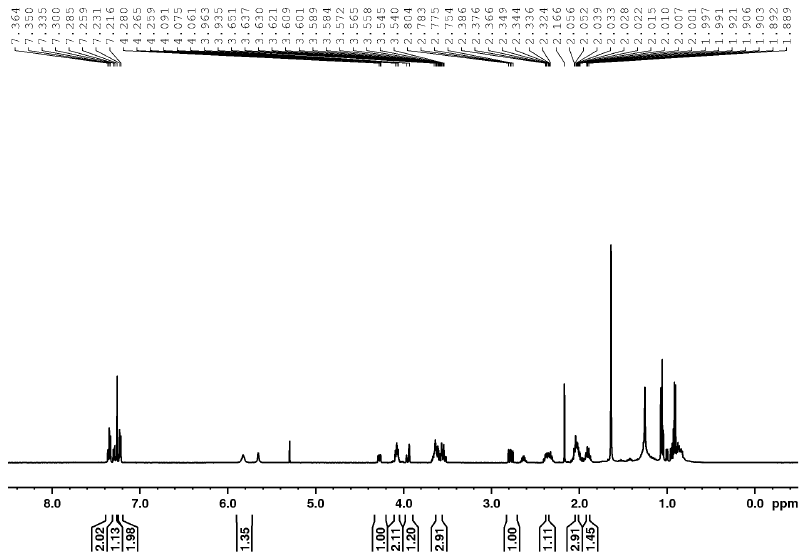

**Figure S2E. ^1^H NMR spectrum of DKP-3 (500 MHz; CDCl_3_)**

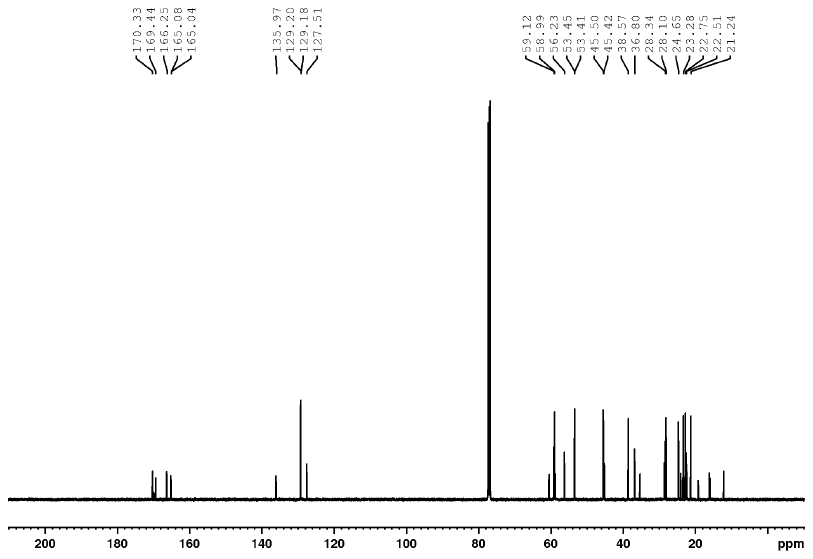

**Figure S2F. ^1^C NMR spectrum of DKP-3 (500 MHz; CDCl_3_)**

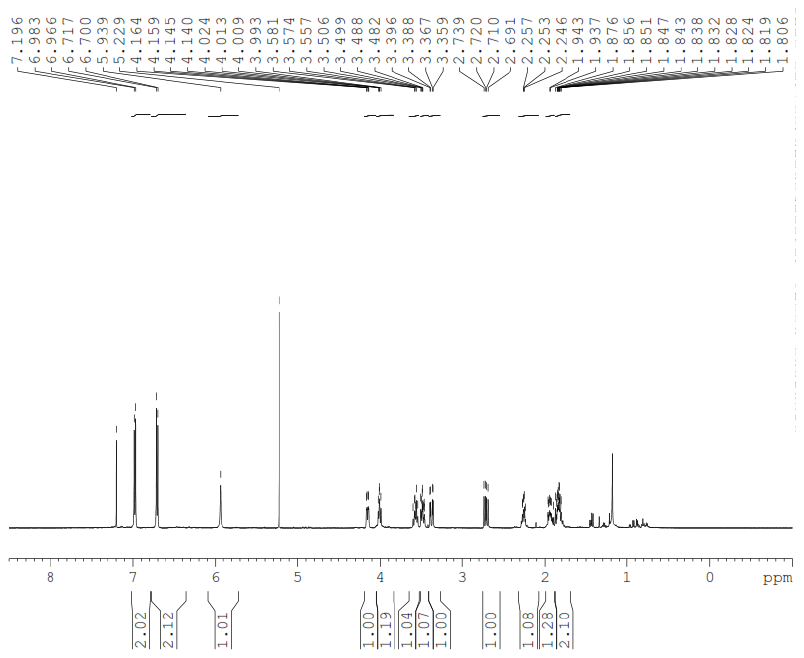

**Figure S2G. ^1^H NMR spectrum of DKP-4 (500 MHz; CDCl_3_)**

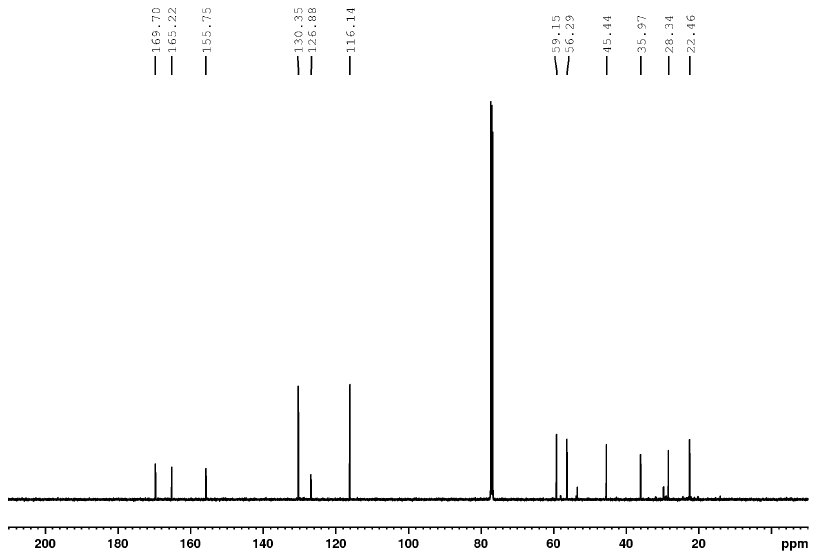

**Figure S2H. ^1^C NMR spectrum of DKP-4 (500 MHz; CDCl_3_)**

**HR-ESI-MS spectra of identified cyclic dipeptides from** *A. veronii* V03

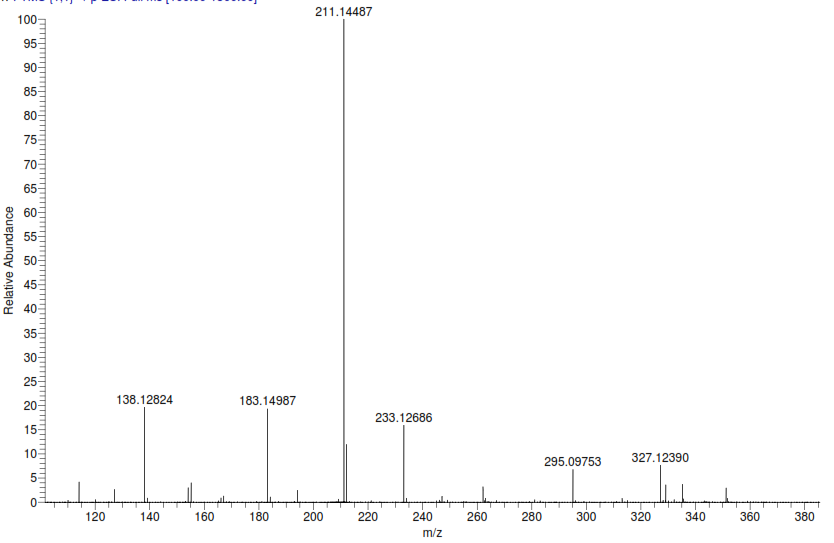

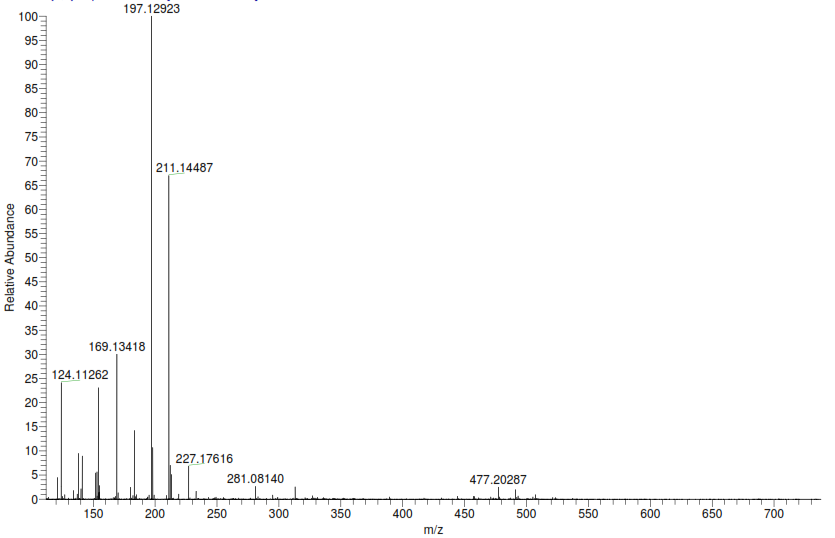
**Figure S3A. Mass spectrum (HRMS) of DKP-1 (M+1)**

**Figure S3B. Mass spectrum (HRMS) of DKP-2 (M+1)**

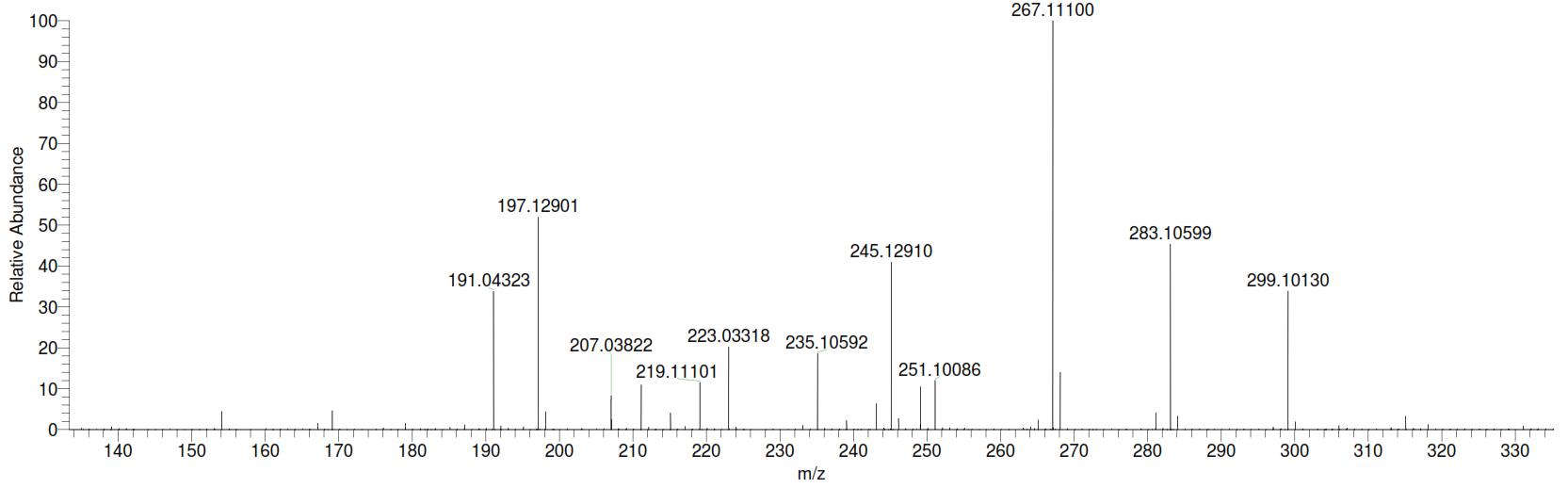

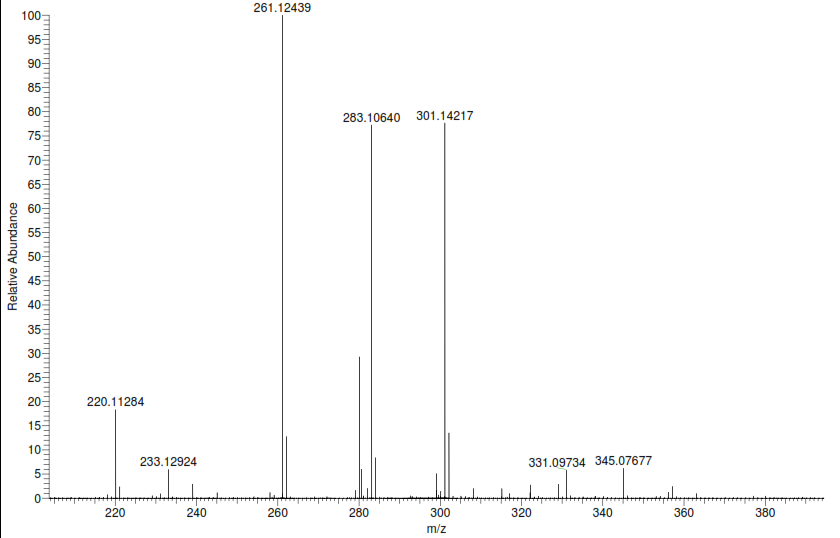

**Figure S3C. Mass spectrum (HRMS) of DKP-3 (M+1)**

**Figure S3D. Mass spectrum (HRMS) of DKP-4 (M+1)**
